# Ferritin nanoparticle based SARS-CoV-2 RBD vaccine induces persistent antibody response and long-term memory in mice

**DOI:** 10.1101/2020.12.22.423894

**Authors:** Wenjun Wang, Baoying Huang, Yanping Zhu, Wenjie Tan, Mingzhao Zhu

## Abstract

Since the outbreak of COVID-19, over 200 vaccine candidates have been documented and some of them have advanced to clinical trials with encouraging results. However, the antibody persistence over 3 months post immunization and the long-term memory have been rarely reported. Here, we report that a ferritin nanoparticle based SARS-CoV-2 RBD vaccine induced in mice an efficient antibody response which lasts for at least 7 months post immunization. Significantly higher number of memory B cells were maintained and a significantly higher level of recall response was induced upon antigen challenge. Thus, we believe our current study provide the first information about the long-term antibody persistence and memory response of a COVID-19 vaccine. This information would be also timely useful for the development and evaluation of other vaccines.

## INTRODUCTION

Since the reported outbreak of severe acute respiratory syndrome coronavirus 2 (SARS-CoV-2) infection in December 2019, SARS-CoV-2 has quickly spread over the world. To date, more than 70 million infection cases have been reported with over 1.6 million deaths (World Health Organization). So far, there is still no effective treatments available. A safe and effective vaccine is highly demanded ^1^.

For an effective vaccine, antibody persistence and long-term memory are favorable features ^2^. The poor antibody persistence during natural SARS-CoV-2 infection raised concerns whether a vaccine could induce a long-lasting antibody response and whether memory recall response would be induced upon reinfection ^3,4^. Currently, over 200 vaccine candidates have been documented and some of them have advanced to clinical trials with encouraging results ^5^. However, to our knowledge, none of them has reported the antibody persistence over 3 months post immunization, and the long-term memory is also unclear ^6–18^. Here, we report that a ferritin nanoparticle based SARS-CoV-2 RBD vaccine induced in mice an efficient antibody response which lasts for at least 7 months post immunization. Significantly higher number of memory B cells were maintained and a significantly higher level of recall response was induced upon antigen challenge.

## RESULTS AND DISCUSSION

### Molecular design and characterization of Ferritin-NP-RBD vaccine

SpyTag/SpyCatcher technique based click vaccine platform has been developed and widely used in our lab, which achieves rapid and convenient production of nanoparticle vaccines ^19,20^. The same strategy was applied for the construction of ferritin NP based SARS-CoV-2 receptor binding domain (RBD) vaccine (Fig. 1a). SpyTag was genetically fused to the C-terminus of SARS-CoV-2 S RBD and the fusion protein was expressed in 293F. Due to glycosylation modification, the apparent molecular weight of RBD-SpyTag was about 35kDa (actual molecular mass was 27.4 kDa). Purified RBD-SpyTag was mixed with SpyCatcher-ferritin NP at different ratios. SpyTag and SpyCatcher mediated covalent conjugation was confirmed by SDS-PAGE. The ferritin-NP-RBD was further purified by size exclusion chromatography. SDS-PAGE analysis confirmed the purity of ferritin-NP-RBD.

**Fig 1.**
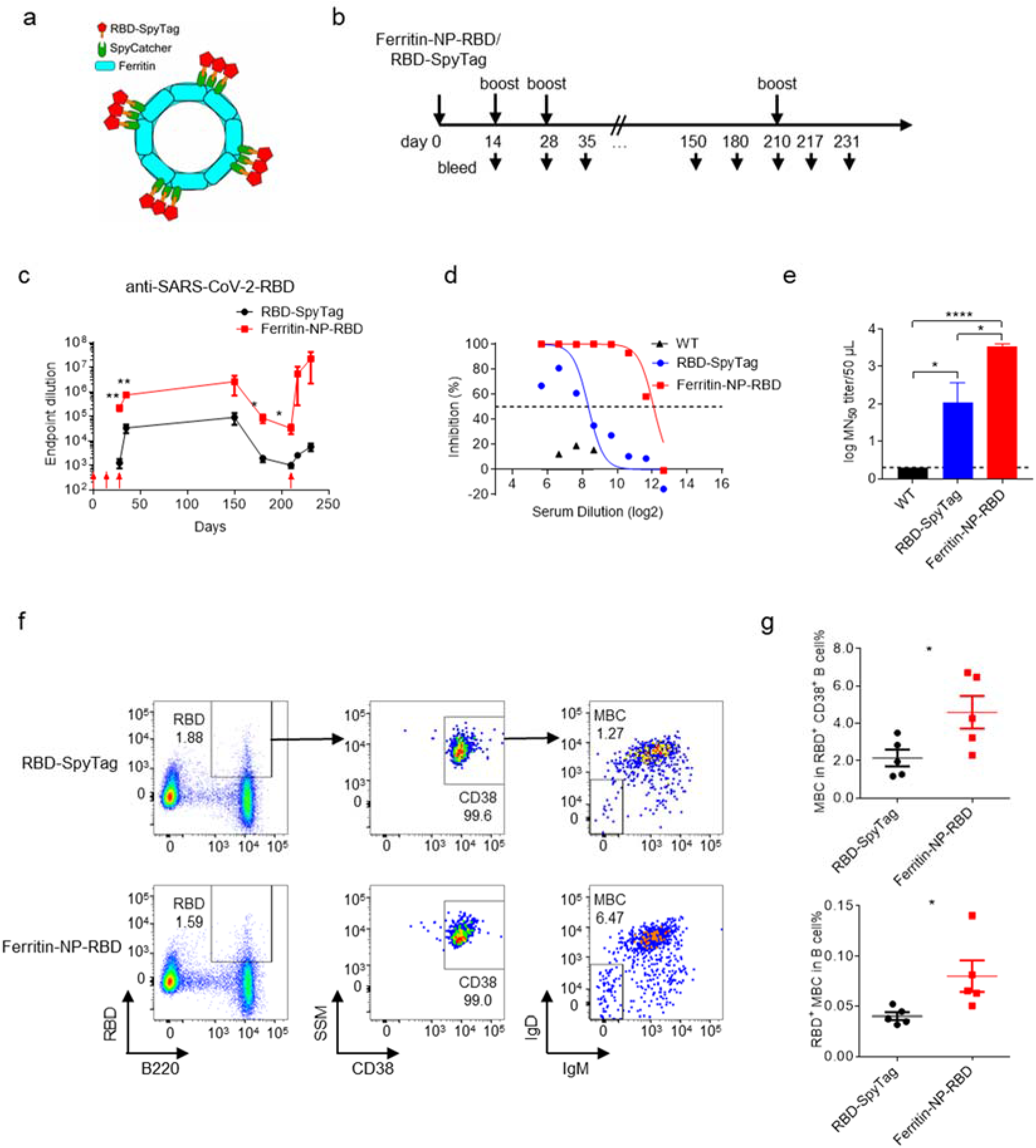
Ferritin-NP-RBD vaccine induces persistent antibody response and long-term memory. **a**, Schematic illustration of ferritin-NP-RBD vaccine construction. **b**, Naïve WT C57BL/6 mice (n=5) were subcutaneously immunized with 500 pmol ferritin-NP-RBD vaccine or equimolar RBD-SpyTag soluble antigen with 30 μg CpG-1826 at day 0, 14 and 28. At 7 months (day 210) after the first immunization, mice were boosted with 200 pmol ferritin-NP-RBD vaccine or equimolar RBD-SpyTag soluble antigen with 30 μg CpG-1826. Blood samples were collected at indicated time points. The red arrows indicate the immunization time points. **c**, Anti-RBD response were monitored and analyzed by ELISA. **d** and **e**, Vero cells were incubated with live SARS-CoV-2 in the presence of immune sera collected from ferritin-NP-RBD or RBD-SpyTag immunized mice at day35 or WT unimmunized mice. Cytopathic effects (CPE) was observed 48 hours post infection. The inhibition (**d**) and MN_50_ titer (**e**) were calculated. **f** and **g**, At 6 months after the first immunization, memory B cell in the peripheral blood was presented (**f**) and statistically analyzed (**g**). Numbers adjacent to the outlined areas indicate percent of each gate. The red arrows indicate the boost immunization time points. Data are shown as mean ± SEM, statistical significance was determined by unpaired two-tailed *t*-test.

### Ferritin-NP-RBD vaccine induces efficient antibody response

To assess the immunogenicity of the ferritin-NP-RBD, naïve wild type (WT) C57BL/6 mice were immunized with ferritin-NP-RDB vaccine or equimolar RBD-SpyTag as control in the presence of CpG-1826 adjuvant at day 0, 14 and 28 (Fig. 1b). Ferritin-NP-RBD vaccine induced an approximate 100-fold higher antibody level than soluble RBD-SpyTag at day 28 (Fig. 1c). After the third immunization, the control vaccine group reached to about 10^5^ antibody titers at day 35, and ferritin-NP-RBD group reached to about 10^6^ antibody titers (Fig. 1c). Thus, RBD conjugated to ferritin-NP greatly enhanced the immunogenicity of RBD antigen and elicited a dramatically enhanced RBD specific antibody response.

### Protective immunity of ferritin-NP-RBD vaccine against SARS-CoV-2

To test whether RBD antisera induced by the ferritin-NP-RBD vaccine could provide protection against the live SARS-CoV-2 *in vitro*, vero cells were infected with live SARS-CoV-2 (C-Tan-nCoV strain 04) in the presence of day 35 sera from different immunization groups. The results showed that four out of five mice from RBD-SpyTag group neutralized over 50% live-virus at serum dilutions ranged from only 1:100 to 1:400, with average 50% microneutralisation (MN_50_) titer was 10^3.8^/ml (Fig. 1f,g). Strikingly, all of five mice from ferritin-NP-RBD vaccine group had neutralization effect at serum dilutions ranged from 1:1600 to 1:3200, with average MN_50_ of 10^4.8^/ml (Fig. 1f,g). These results confirm that the antisera to RBD elicited by the ferritin-NP-RBD vaccine can prevent SARS-CoV-2 infection much more effectively *in vitro*.

### Ferritin-NP-RBD vaccine induces persistent antibody response and memory response

To determine the antibody persistence induced by ferritin-NP-RBD vaccine in mice, we continued to monitor the antibody responses at 5 months, 6 months and 7 months after the first immunization. The anti-RBD level at 5 months was almost comparable to the level of day 35 (Fig. 1c). At 6 and 7 months, while the antibody endpoint titers of both groups gradually dropped, ferritin-NP-RBD vaccine group still maintained significantly higher level of anti-RBD than the RBD-SpyTag control vaccine group (Fig. 1c), confirming the benefit of ferritin-NP for maintaining antibody response. To further determine whether ferritin-NP promotes better memory response, we first examined the RBD-specific memory B cells (MBCs) in the blood. Significantly more RBD-specific MBCs were generated and maintained at 6 months in ferritin-NP-RBD group compared with RBD-SpyTag control group (Fig. 1f,g). Consistent with the enhanced MBC formation and maintenance, when mice were challenged with RBD vaccine antigen at day 210, ferritin-NP-RBD group elicited a dramatically increased antibody recall response, 2000 times stronger than that in control group (Fig. 1c). Thus, Ferritin-NP-RBD vaccine induced not only a persistent RBD-specific antibody response but also long-term memory.

Self-assembling ferritin NPs have recently become widely used for vaccine design ^19,21–24^. It also emerged as an attractive platform for SARS-CoV-2 vaccine design ^10^. In this study, a similar approach was used for vaccine construction as we used previously and here ^19,20^. Upon two immunizations, about 10^5^ titers of RBD-specific anti-IgG was detected. In our study, an average of 2.2×10^5^ titers of antisera were induced upon two immunizations, and about 10^6^ titers of antisera were induced upon three immunizations. Given such impressive primary antibody response, we have further monitored antibody persistence and memory response throughout 7 months, which is so far the longest reported period for COVID-19 vaccine evaluation, to our knowledge. The extended antibody persistence and well-boosted recall antibody response as demonstrated in current study warrant future success of ferritin-based COVID-19 vaccine.

Currently, multiple SARS-CoV-2 vaccine candidates, such as inactivated virus vaccine^7^, vectored vaccine^11,13,18^, mRNA vaccine^8,9,17^ and protein subunit vaccine^6,12,14,16^, are under development and clinical trials. Although vaccines come in different forms and are immunized in different doses, our ferritin based NP vaccine induced roughly equal antibody titers (endpoint titer 10^6^) and live SARS-CoV-2 neutralizing activity compared with inactivated vaccine PiCoVacc^7^, mRNA based vaccines^9,17^ and RBD-sc-dimer protein subunit vaccine^6^. In addition, more importantly, current ferritin-NP-RBD vaccine has demonstrated persistent antibody response and impressive long-term memory.

## Methods

### Mice

Naïve WT C57BL/6 mice were obtained from SPF (Beijing) Biotechnology Co.,Ltd.. Mice were housed under specific pathogen-free conditions in the animal care facilities at the Institute of Biophysics, Chinese Academy of Sciences. All animal experiments were performed in accordance with the guidelines of the Institute of Biophysics, Chinese Academy of Sciences, using protocols approved by the Institutional Laboratory Animal Care and Use Committee.

### Cloning, expression, and purification of fusion proteins

The SpyCatcher-ferritin nanoparticle vaccine platform was prepared as described previously ^20^. Briefly, pDEST14-SC-(G4S)_3_-ferritin plasmid was expressed in BL21 (DE3) competent E. coli cells, and purified with superpose 6 increase (GE Healthcare Life Sciences, Pittsburgh, PA, USA) size exclusion column.

RBD-SpyTag was expressed and purified from 293F, 6 His-tagged RBD-SpyTag were cloned into pEE12.4 vector. pEE12.4-RBD-SpyTag-Histag plasmids were transfected with PEI (PEI MAX-Transfection Grade LinearPolyethylenimine Hydrochloride, Polysciences 24765-1) into 293F cells. 24 hours after transfection, 3.8 mM of final concentration VPA (valproic acid, sigma P4543) was added into culture to inhibit cell growth, then incubated for about 6 days before the final collection. Supernatants were collected and centrifuged at 10000 rpm, 4°C。Discard the cellular debris and incubated the supernatants with Ni-NTA agarose to enrich RBD-SpyTag protein, followed by elution with PBS buffer containing 100 mM imidazole. The purified proteins were concentrated and buffer-replaced with PBS. The target RBD-SpyTag protein were confirmed by SDS-PAGE and size exclusion chromatography.

### Generation and purification of ferritin-NP-RBD vaccine

The purified RBD-SpyTag was conjugated to SC-ferritin-NP *in vitro* to construct the ferritin-NP-RBD vaccine. To assay reconstitution, SC-ferritin-NP was reacted with RBD-SpyTag at molar ratio of 1:0.5, 1:1, and 1:2 at 4 °C overnight. SDS-PAGE was used to evaluate the reconstitution efficiency. For vaccine generation, SC-ferritin-NP was mixed with RBD-SpyTag at a molar ratio of 1:1.5 at 4 °C overnight. The conjugated ferritin-NP-RBD was then purified by a Superose 6 Increase size exclusion column (Ve=12ml, Vt= 24 ml, Vo= 8 ml).

### Immunization

Female naïve WT C57BL/6 mice (8-9 weeks old) were subcutaneously immunized with 500 pmol (approximately 30.7 μg, as determined by single ferritin-RBD subunit) ferritin-NP-RBD vaccine or 500 pmol (approximately 13.7 μg) RBD-SpyTag with 30 μg CpG-1826 (Generay Technology, Shanghai, China), respectively, at the tail base at day 0, day 14 and day 28. For the memory response assay, another boost immunization was performed at day 210. 200 pmol (approximately 12.3 μg) ferritin-NP-RBD vaccine or 200 pmol (approximately 5.5 μg) RBD-SpyTag with 30 μg CpG-1826, respectively, was used.

### ELISA

For the RBD-specific ELISA, 5 μg/ml RBD-SpyTag protein produced in our lab was coated onto 96-well high binding Costar^®^ Assay plates (CORNING) at 4 °C overnight. After blocking with a blocking buffer (PBS containing 5% FBS), serum samples with different dilutions were added onto the plates. Horseradish peroxidase (HRP) conjugated goat anti-mouse IgG (H+L) (1:5000, ZSGB-BIO) was used as second antibody. The concentration of specific antibodies was measured using TMB substrate (SeraCare) and the absorbance at 450 nm-630 nm was detected by a Microplate Reader (Molecular Devices).

### Live SARS-CoV-2 neutralization assay

The experiment was conducted in a BSL-3 laboratory, as previously reported ^25–28^. Briefly, the sera were 2-fold serially diluted using 2% FBS-DMEM and mixed with the same volume of live SARS-CoV-2 (C-Tan-nCoV strain 04, 100TCID50), the mixtures were incubated at 37 °C for 1 h, following which they were added to the seeded Vero cells. After incubation at 37 °C for 48 h, CPE was observed, and 100 μL of the culture supernatant was harvested for nucleic acid extraction and realtime fluorescence RT-PCR reaction. Median tissue culture infective dose (TCID50) of the virus in the sample was calculated according to the CT value of the sample and standard curve. The neutralization potency (or inhibition rate) was calculated as follows: Inhibition ratio = (TCID50 without serum - TCID50 with serum)/TCID50 without serum *100%. The median micro-neutralization dose (MN_50_) was calculated by Reed-Munch method.

### Flow cytometry

Fresh blood samples were collected from mice at 6 months after the first immunization. Red blood cells were lysed with Ammonium-Chloride-Potassium (ACK) Lysing Buffer. Single cell suspensions were resuspended in an appropriate volume of FACS buffer (1-5×10^6^ cells/100 μl), blocked with anti-FcγR mAb (clone 2.4G2) to block nonspecific binding. For RBD-specific memory B cells (MBC) staining, biotin conjugated recombinant RBD protein was incubated with cell suspensions together with fluorescence-conjugated antibodies, then RDB binding cells were recognized by APC/Cy7 Streptavidin (Biolegend). The antibodies used included anti-B220-PE (RA3-6B2, eBioscience), anti-CD38-APC (90, eBioscience), anti-IgM-PE/Cy7 (II/41, eBioscience) and anti-IgD-PerCP/Cy5.5 (11-26c.2a, Biolegend). The working concentration of the antibodies is 2.5μg/ml.

### Statistical analysis

All analysis was performed using GraphPad Prism statistical software (GraphPad Software Inc., San Diego, CA, USA). All of the data were analyzed using unpaired two-tailed *t*-test. The results are expressed as the mean ± SEM. A value of P<0.05 was considered statistically significant.

## Acknowledgements

This work was supported by grants from Strategic Priority Research Program of the Chinese Academy of Sciences (XDB29040202 to M.Z.), National Key R&D Program of China (2019YFA0905903 to M.Z.).

## Author contributions

W.W. conducted vaccine preparation, immunization, antibody titering and memory B cell determination; B.H. conducted anti-sera neutralizing assay; Y.Z. prepared vaccines; W.W., B.H., Y.Z., W.T. and M.Z designed the experiments, analyzed the data, and wrote the manuscript; W.T. and M.Z supervised the project; M.Z conceived the project.

